# Phytochemical screening, antioxidant properties and antihyperlipidemic activity of *Chrysobalanus icaco* L. seeds

**DOI:** 10.1101/2025.09.12.675981

**Authors:** Jacinta E. Apitikori-Owumi, Mubo A. Sonibare

## Abstract

*Chrysobalanus icaco* L. (Chrysobalanaceae) is a widely utilized local spice prevalent in the coastal region of the Niger Delta. *C. icaco* seeds are traditionally documented for their therapeutic applications in treating various conditions, including chronic hyperlipidemia, diabetes, and hemorrhage. This study aimed to investigate the phytochemical, antioxidant, and antihyperlipidemic properties of *C. icaco* seed extract. Phytochemical analysis identified 16 secondary metabolites in the crude extract. Extract of *C. icaco* demonstrated a reduction in total cholesterol, triglycerides, low-density lipoproteins (LDL), and very low-density lipoproteins (VLDL) levels, and an increase in high-density lipoprotein (HDL) level within the treatment groups relative to the negative control group. Anti-hyperlipidemic activity confirmed the traditional use of *C. icaco* in lowering cholesterol levels.

## INTRODUCTION

Hyperlipidemia is a significant public health concern globally (Bencheikh et al., 2021) and a chronic disorder of metabolic syndrome characterized by an elevated amount of lipids. It accounts for 17 million deaths worldwide annually (Addisu et al., 2023; Kubi et al., 2024). Hyperlipidemia is a significant risk factor for cardiovascular and cerebrovascular diseases, which are the leading causes of death and disability globally, and eighty percent of cardiovascular deaths occur in developing countries (Stanaway et al., 2018; Hussain et al., 2024).

Recently, interest in the search for new antihyperlipidemic agents from natural products with lower side effects, safety, therapeutic impact, and local availability has increased. Antihyperlipidemic natural products promote a reduction in cholesterol levels in the blood (Asija et al., 2016, 2022; Al-Snafi, 2022).

*Chrysobalanus icaco*, widely referred to as cocoplum, is a member of the Chrysobalanaceae family. *C. icaco* is well-regarded among the Ijaw people in the villages of Ezetu and Kolomao in Bayelsa State, southern Nigeria, where it is commonly referred to as ‘Ebulo’ by the Izon and ‘Omilo’ by the Itsekiri (Apitkori-Owumi et al., 2024). It grows naturally along the sea coast in the tropical regions of America, Central America, and Africa (De Aguiara et al., 2017; Silver et al., 2017; Hanish et al., 2020). This spice is widely used by the Itsekiri, Ukwauni, Urhobo, and Izon tribes in the Niger Delta region of Nigeria to prepare pepper soup, a cherished delicacy. In various countries, including Mexico, El Salvador, the United States, Trinidad, and Brazil, *C. icaco* has been employed in traditional medicine to address high cholesterol, diabetes, chronic diarrhea, bleeding, abdominal pain, dysentery, leucorrhoea, anemia, and inflammation. Previous studies have demonstrated the beneficial effects of aqueous *C. icaco* in treating obesity, enhancing glycemic homeostasis, and preventing fat storage in Wistar rats (Portela-de-Sá et al., 2020). Previous research reported that *C. icaco* leaves can be used to treat various ailments, particularly obesity, but there is a pressing need to explore the use of *C. icaco* seeds as an antihyperlipidemic agent, as no data currently exists on the antihyperlipidemic properties of *C. icaco* seeds. Consequently, this study was conducted to investigate the antihyperlipidemic properties of *C. icaco* seeds.

## MATERIALS AND METHODS

### Collection, authentication and extraction of plant

Viable dried seeds of *C. icaco* were procured from the principal market in Warri, Delta State, Nigeria. The identification and authentication of the fruits were conducted at the Department of Botany and Biotechnology, University of Benin, Benin City, and were assigned the voucher number UBH-C393. The seeds of *C. icaco* were pulverized using an electric blender. The sample underwent extraction with 70% ethanol employing a Soxhlet extractor. The extract was subsequently concentrated using rotary vacuum evaporation.

### Qualitative phytochemical screening

The phytochemical analysis was carried out using the approach detailed by Silva et al. (2017); Singh and Kumar (2017).

### Quantitative analysis of phytochemicals in the seed extract

The phytochemical composition of the seed extract was quantitatively analysed by using procedures established by Mallikarjuna et al. (2012) and Saleem et al. (2018).

### Evaluation of antioxidant activity

The free radical scavenging activity (2, 2-diphenyl-1-picryl-hydrazyl) was measured using a protocol established by Karak (2019), and a ferric reducing antioxidant power (FRAP) assay was evaluated according to the method proposed by Chandha and Dave (2009).

### Animal care and experimental design

The study received ethical approval from Delta State University’s College of Health Sciences’ Ethical Review Committee in Abraka, Nigeria. The number for the ethical permit was REC/FBMS/DELSU/22/145.The study was carried out in compliance with globally recognized guidelines for laboratory animal care and care outlined in the European Community Guidelines (1986 Directives 86/609/E. E. C) as well as the Guidelines for the United States and Canada (NRC. I. H. Publication 8523; 1985 revision).

The rats were acclimated in the Animal House of the Faculty of Basic Medical Sciences, College of Health Sciences, Delta State University. The standard rat feed used to feed the animals was acquired from Maiestas Herbagro Enterprise in Abraka, Delta State, Nigeria. The animals were kept in constant water throughout the study. Throughout the study, male and female Wistar rats were housed in different animal cages to avoid mating and reproduction (Asuzu-Samuel and Apitikori-Owumi, 2023). A test for acute toxicity was performed using the Lorke method (Chinedu et al., 2013).

Following acclimatization, the animals were randomly assigned into eight groups (n=10): animals that receive standard diet only without hyperlipidemia induction (group 1), animals induced with hyperlipidemic without treatment (group 2), induced hyperlipidemic animals treated with atorvastatin (2.0 mg/kg body weight) (group3) and five treatment groups for the seed extract (100, 200, 300, 400 and 500 mg/kg body weight) (groups 4 to 8).

Hyperlipidemia was induced in the animals using the method described by Nwankwo et al. (2021) with slight changes. A hundred and fifty grams of a high-fat diet was administered orally to the animals for 28 days (4 weeks), except for those in group 1. Doses of seed extract and standard drug were administered orally to the animals. Weights of the animals were measured and documented weekly. Following the twenty-eight days of inducing hyperlipidemia, five animals each from groups 1 and 2 were sacrificed using the technique described by Akinlolu et al. (2020). Blood was collected via cardiac puncture from animals from animals for haematological profile and biochemical profile (total cholesterol, triglyceride, low-density lipoprotein (LDL) cholesterol, high-density lipoprotein (HDL) cholesterol, aspartate aminotransferase (AST), alkaline phosphatase (ALP), alanine aminotransferase (ALT), urea, and creatinine).

The procedure for establishing hyperlipidaemia in group 2 involved comparing their lipid levels with those of group 1, successfully inducing the condition. This procedure certified hyperlipidaemia in group 2. Subsequently, the animals were administered 100, 200, 300, 400, and 500 mg/kg body weight of the extract or atorvastatin over a period of twenty-eight days (4 weeks). Group 2, however, received no treatment and was provided with standard animal feed and water. Upon completion of the experiment, all animals were euthanized utilizing the technique specified by Akinlolu et al. (2020).

Blood samples were collected, and the biochemical and haematological profiles were analyzed as previously described. The liver, heart, and kidneys from the animals underwent a histopathological investigation.

### Statistical analysis

The data collected from the quantitative analysis of phytochemicals, antioxidants, animal lipid profiles, liver enzymes, creatinine and urea levels, and haematological profiles were statistically evaluated using one-way analysis of variance (ANOVA) in SPSS version 26.0 (IBM Corp., U.S.A.) software for Windows. A P-value of ≤ 0.05 was statistically considered significant.

## RESULTS

### Qualitative phytochemical screening of seed extract of *C. icaco*

The analysis indicated that the seed extract of *C. icaco* is rich in various phytochemicals, including alkaloids, tannins, phenolic compounds, flavonoids, cholesterol, steroids, terpenoids, triterpenoids, phytosterols, saponins, cardiac glycosides, carbohydrates, reducing sugars, proteins, amino acids, fats, and oils (Table 1).

**Table 1.** Qualitative phytochemical screening of seed extract of *C. icaco*.

| Phytochemicals | Ethanol extract |
| --- | --- |
| Alkaloid | +++ |
| Tannin | +++ |
| Phenolic compounds | ++ |
| Flavonoid | ++ |
| Cholesterol | + |
| Terpenoid | + |
| Triterpenoids | + |
| Phytosterol | ++ |
| Saponin | +++ |
| Cardiac glycoside | - |
| Carbohydrate | ++ |
| Reducing sugar | ++ |
| Protein | + |
| Amino acid | + |
| Fats and oil | +++ |

### Quantitative phytochemical screening of seed extract of *C. icaco*

The results indicated that the tannin content was the most significant at 12.48±0.01, followed by phenolic compounds content at 11.63±0.03, alkaloids at 5.50±0.03, and flavonoids content at 2.35±0.06 (Table 2).

**Table 2.** Phytochemical content of the ethanol extract of *C. icaco* seeds.

| Chemical constituents | Standard (mg Gallic Acid Equivalents-dry matter) |
| --- | --- |
| Total tannin | 12.48±0.01 |
| Total phenol | 11.63±0.03 |
| Chemical constituent | Standard (mg Quercetin Equivalents /g dry matter) |
| Total flavonoid | 2.35±0.06 |
| Chemical constituent | Standrd (AE/g dry matter) |
| Total alkaloid | 5.50±0.03 |

### Antioxidant activities of seed extract of *C. icaco*

The findings demonstrated that antioxidant activities increased in a dose-dependent manner. At a concentration of 50 µg/mL), the DPPH activity was 25.81± 6.75, while the FRAP value was 18.47 ± 2.7. At a concentration of 100 µg/mL), the DPPH activity was 35.35 ± 2.57, and (Table 3).

**Table 3.** Antioxidant activities of the ethanol extract of *C. icaco* seeds.

| Extract | Conc. 50 (µg/mL) | Conc. 100 (µg/mL) | Vitamin |
| --- | --- | --- | --- |
| DPPH | 25.8±6.75 | 35.4±2.57 | 45.9±2.93 |
| FRAP | 18.5±2.70 | 22.1±1.71 | 33.0±1.33 |

### Acute toxicity test of the seed extract of *C. icaco*

The results of the acute toxicity tests of *C. icaco* are presented in Table 4. The experiment was carried out in two phases; phase I animals were divided into three groups (n=3), while phase II animals were divided into three groups (n =1). The ethanol extract from *C. icaco* seeds caused acute toxicity, with an LD50 value of 3807.89 mg/kg.

**Table 4.** Phase I and II acute toxicity test of the seed extract of *C. icaco*.

| Experiment | Dose (mg/kg bw) | No. of dead rats after 24 h | No. of dead rats after 24 h | No. of dead rats after 48 h |
| --- | --- | --- | --- | --- |
| Phase I | 10 | 0/3 | 0/3 |  |
|  | 100 | 0/3 | 0/3 |  |
|  | 1000 | 0/3 | 0/3 |  |
| Control | 0 | 0/3 | 0/3 |  |
| Phase II | 1600 | 0/3 | 0/3 |  |
|  | 2900 | 0/3 | 0/3 |  |
|  | 5000 | 0/3 | 0/3 | 2/3 |

### Weights of hyperlipidaemic animals before treatment with seed extract

Figure 1 illustrates the weights of hyperlipidemic animals before treatment with seed extract, while Figure 2 presents the weights of hyperlipidemic animals after treatment with seed extract. Following a four-week induction of hyperlipidaemia, all groups (groups 2 to 8) exhibited significant weight gain before treatment. The percentage increase in weight gain was significantly greater (p < 0.05) in group 2 than in group 1. The percentage increase in weight gain was significantly greater (p < 0.05) in group 2 compared to group 1. On the other hand, after administering the extract and atorvastatin for 28 days, there was a significant reduction (p < 0.05) in body weight across all treated groups (groups 4 to 8) when compared to group 2 and group 3. Figures 3, 4, 5, and 6 illustrate the outcomes related to lipid profiles, kidney parameters, and liver profiles, respectively. A notable decrease (p < 0.05) was observed in the levels of cholesterol, triglycerides, LDL cholesterol, and VLDL cholesterol, alongside an increase in HDL levels in the treated groups when compared to group 1.

**Figure 1.**
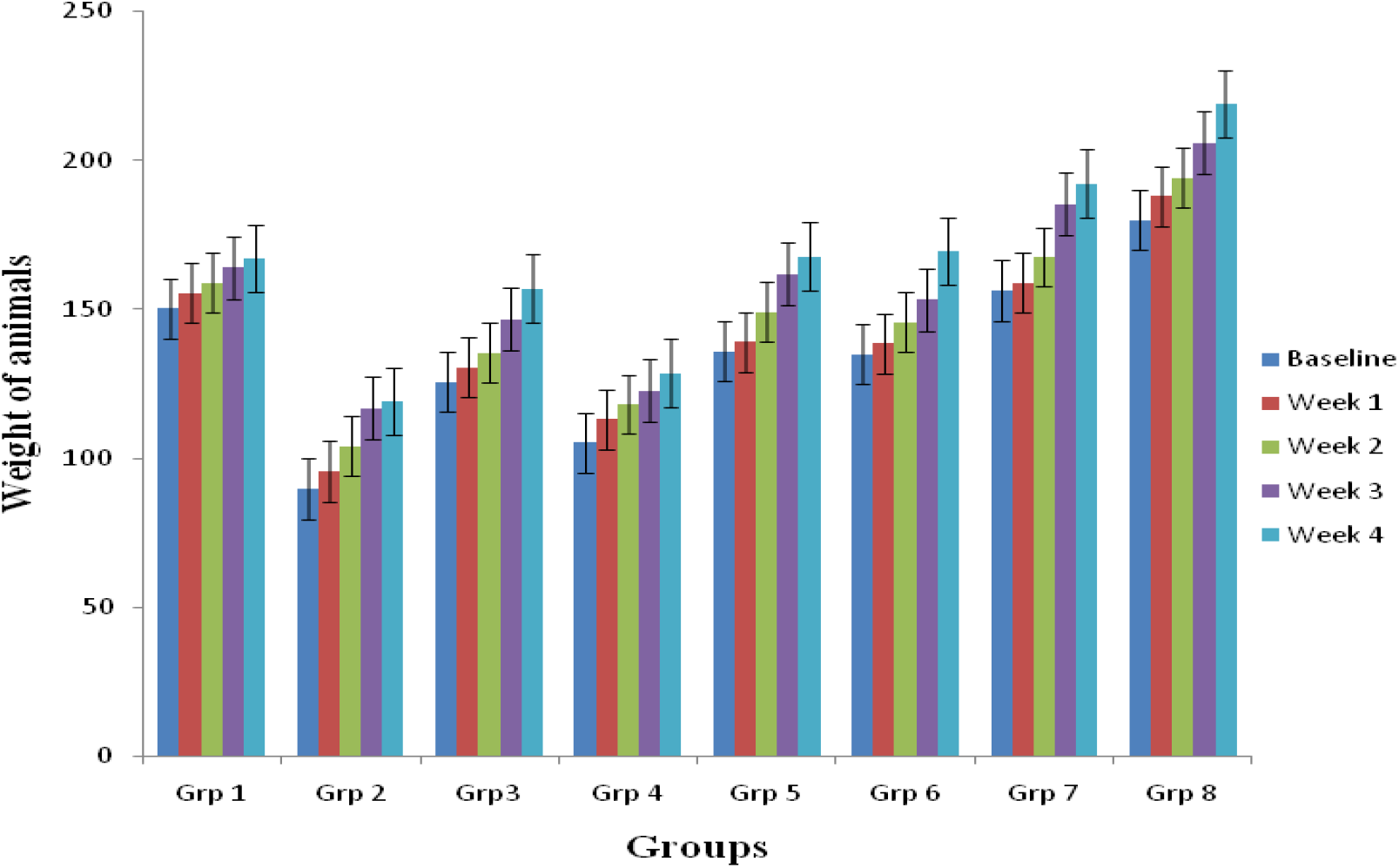
The weights of hyperlipidaemic animals before treatment with the seed extract. The significance level was *p* <0.05. The values are expressed as the mean ± standard error of the mean (SEM), n=10.

**Figure 2.**
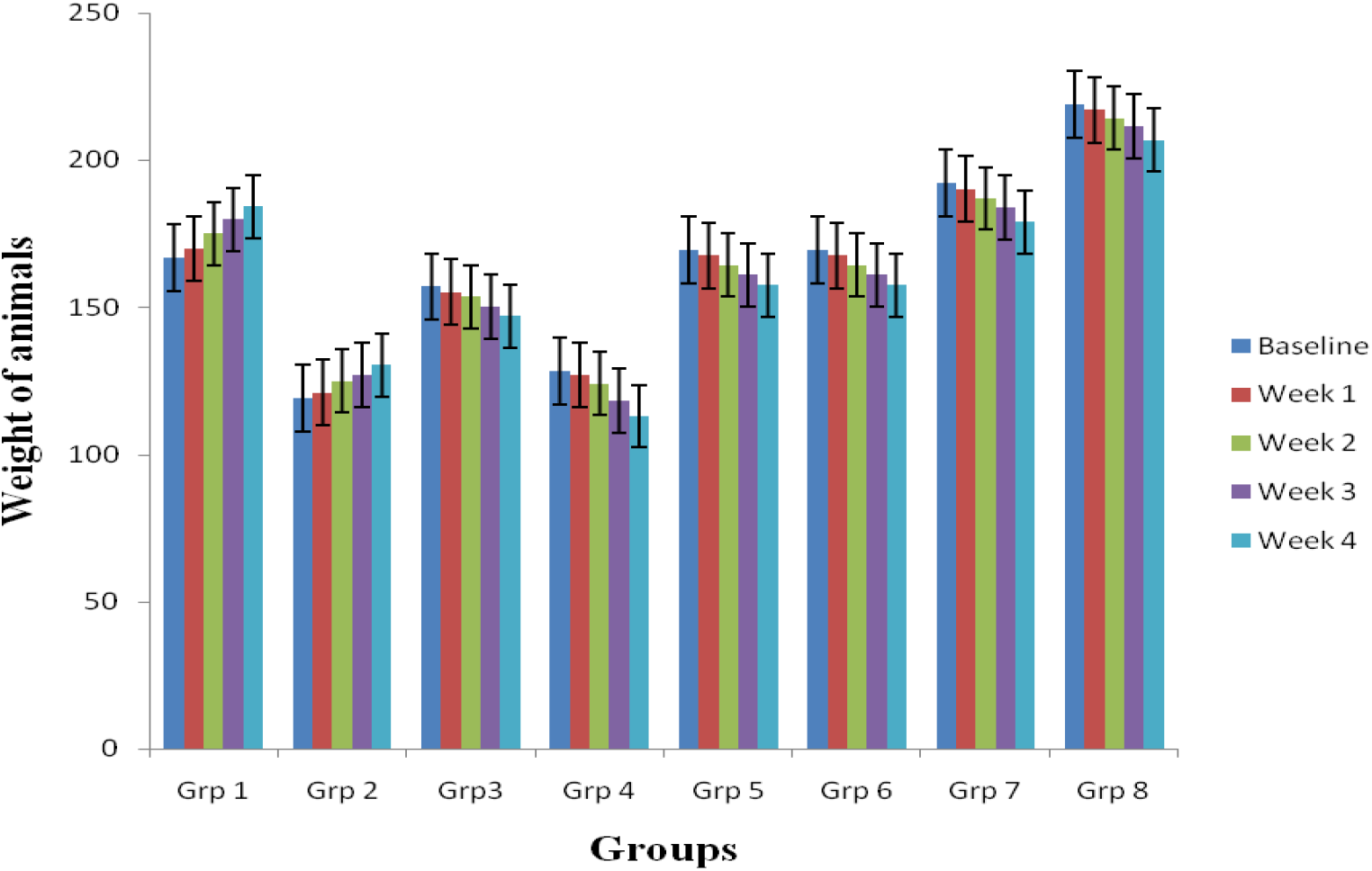
The weights of hyperlipidemic animals after treatment with seed extract. The significance level was *p* < 0.05. The values are expressed as the mean ± standard error of the mean (SEM), n=10.

**Figure 3.**
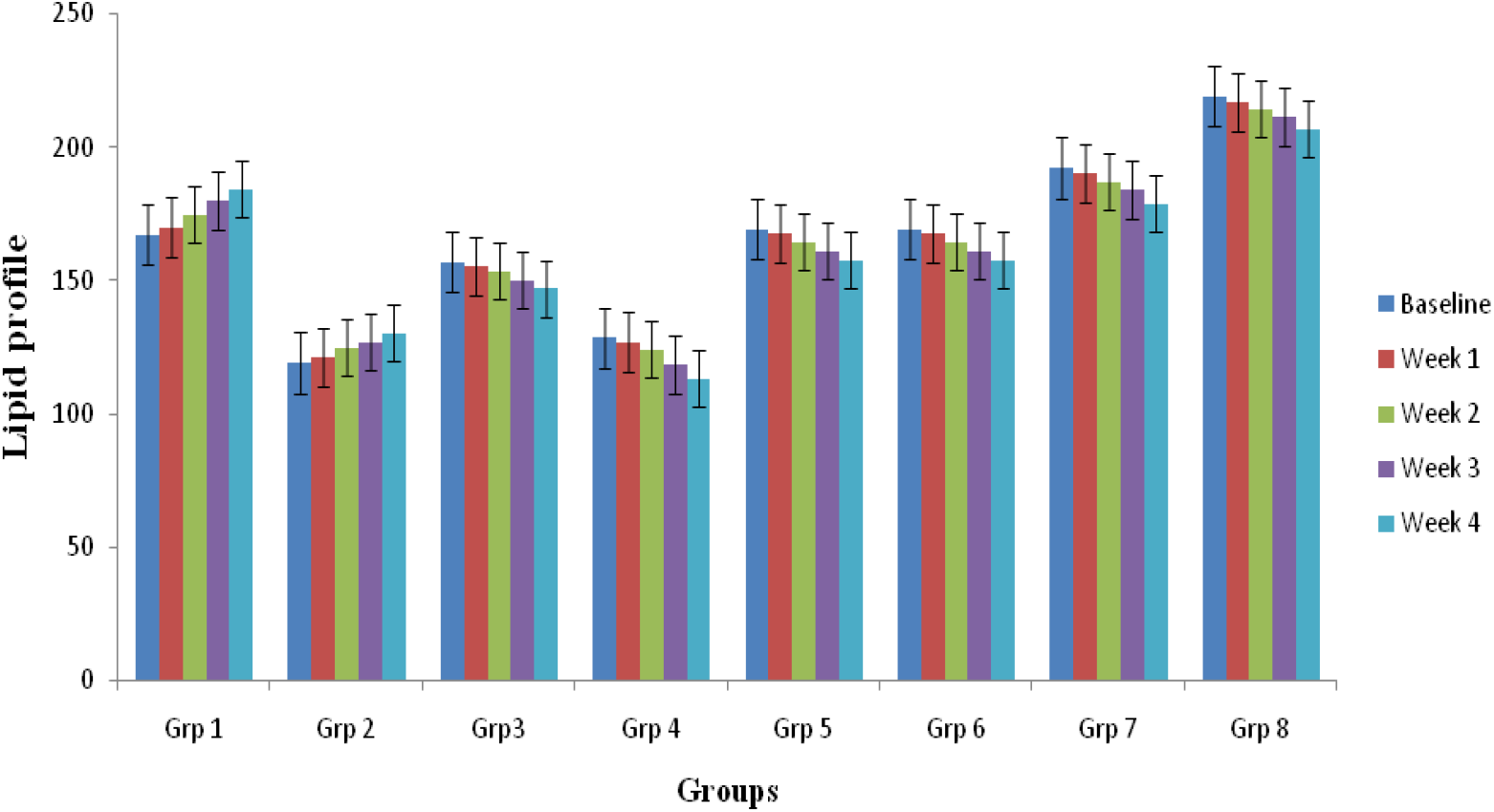
Lipid profile of hyperlipidaemic animals treated with seed extract. The significance level was *p* < 0.05. The values are expressed as the mean ± standard error of the mean (SEM), n=10.

**Figure 4.**
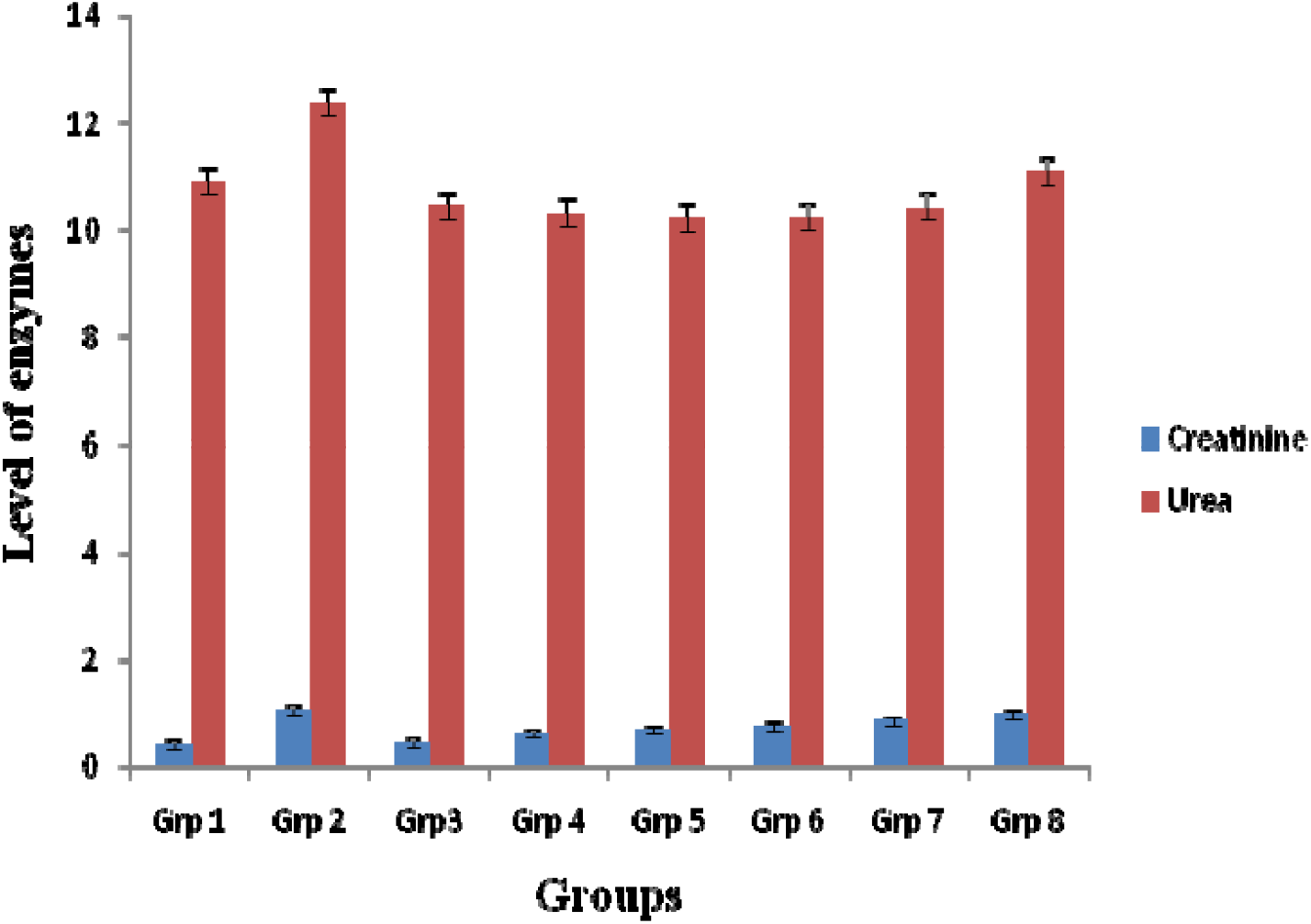
The creatinine and urea profiles of hyperlipidaemic animals treated with seed extract. The significance level was *p* < 0.05. The values are expressed as the mean ± standard error of the mean (SEM), n=10.

**Figure 5.**
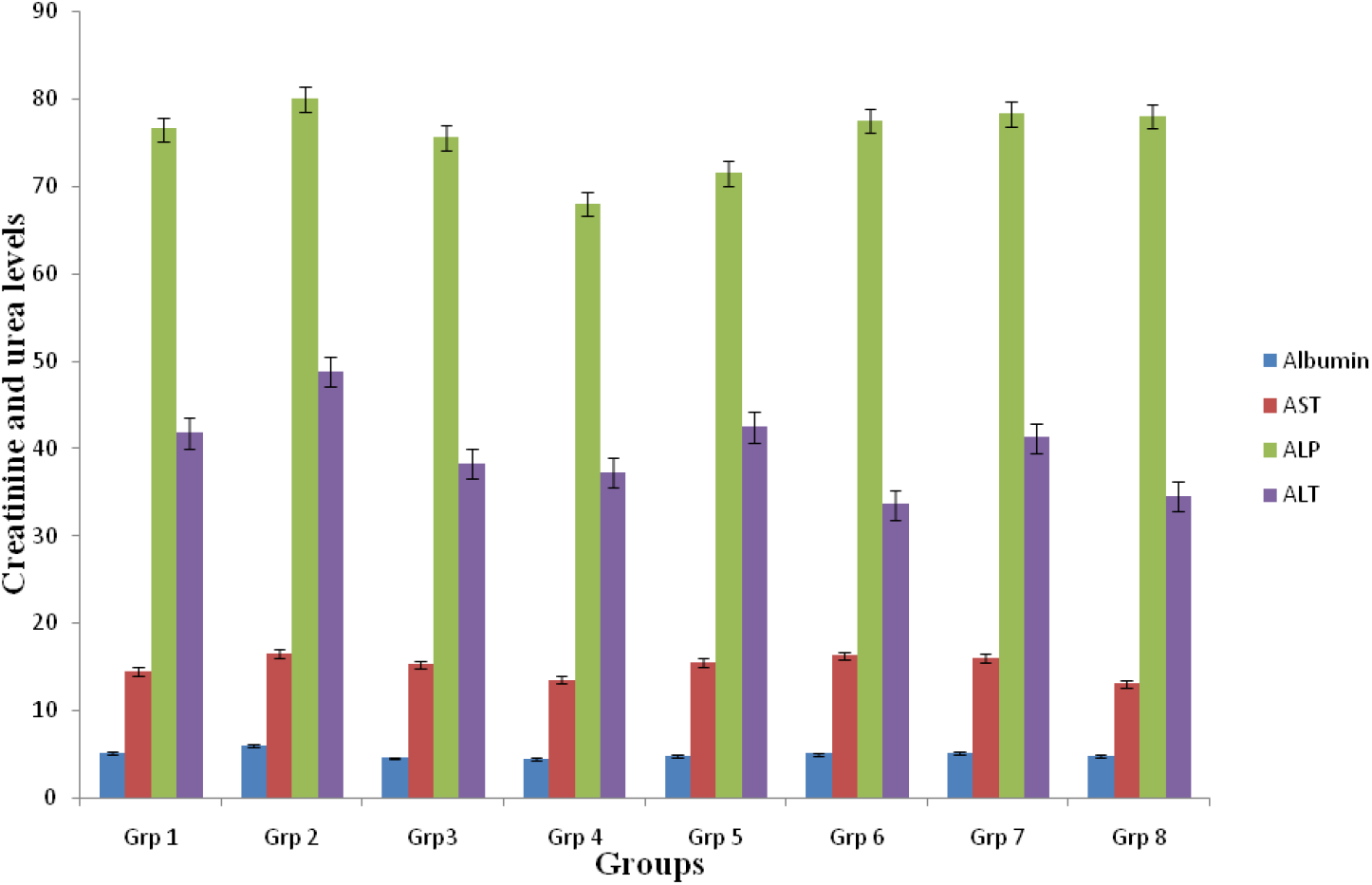
Liver profile of hyperlipidaemic animals treated with ethanol seed extract. The result is significant at *p* < 0.05. The values are expressed as the means ± standard errors of the means (SEM), n=10.

**Figure 6.**
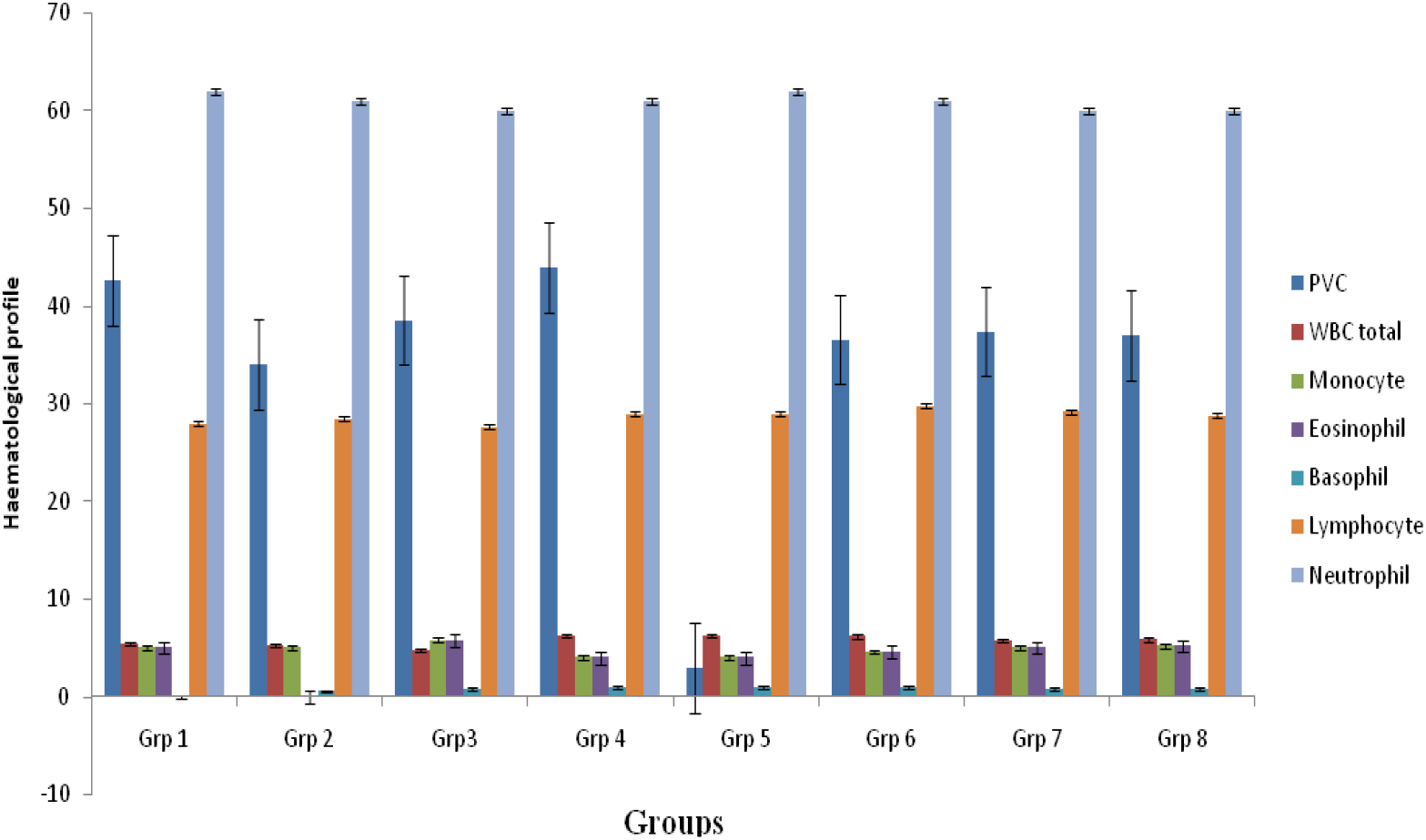
Haematological profile of hyperlipidaemic animals treated with seed extract. The significance level was p <0.05. The values are expressed as the mean ± standard error of the mean (SEM), n=10.

The results indicated that a comparison of groups 3 to 8 with group 2 yielded no statistically significant changes. However, treatment with the extract at doses of 100 mg/kg (group 4) and 200 mg/kg (group 5) led to a significant reduction in ALP values. Additionally, there was a notable decrease in ALT values in groups 3 to 8 when compared to group 2. No significant difference (p < 0.05) was observed in the creatinine or urea levels between groups 3 to 8 and group 2.

Following the administration of the extract alongside the standard drug, there was no significant variation in haematological parameters (WBC total, eosinophil, neutrophil, monocyte, and lymphocyte counts) in groups 3 through 8 relative to groups 1 and 2.

The histological findings for Plates 1-8 are detailed. Following a 28-day treatment period, histological evaluation revealed mild hepatocellular necrosis and inflammation in the negative control group (group 2). The extract’s effects, as demonstrated in Plates 6, 7, and 8, encompassed mild portal inflammation, multifocal hepatocellular necrosis, and hyperplasia of Kupffer cells in the liver of the groups receiving 300, 400, and 500 mg/kg extracts. Following the administration of the extract at concentrations of 300, 400, and 500 mg/kg, as shown in (Plates 6, 7, and 8), inflammation and moderate atrophy of the glomerular tuft were observed in the kidney.

**Plate 1.**
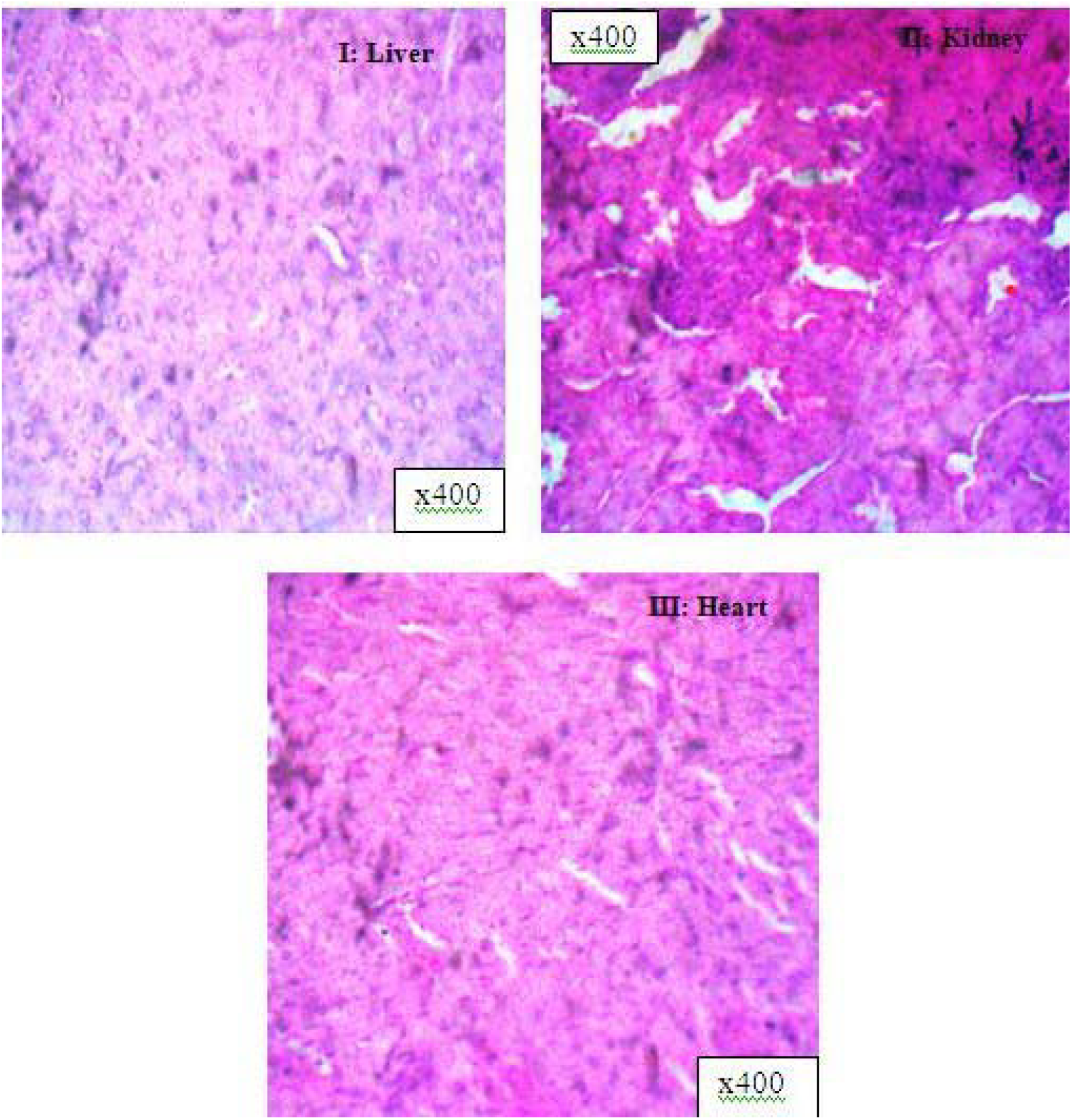
Histology of group 1 animals (normal rats) stained with H&E. i) Liver-There is no observable lesion, ii) Kidney-There is no observable lesion, and III) Heart-There is no observable lesion (x400).

**Plate 2.**
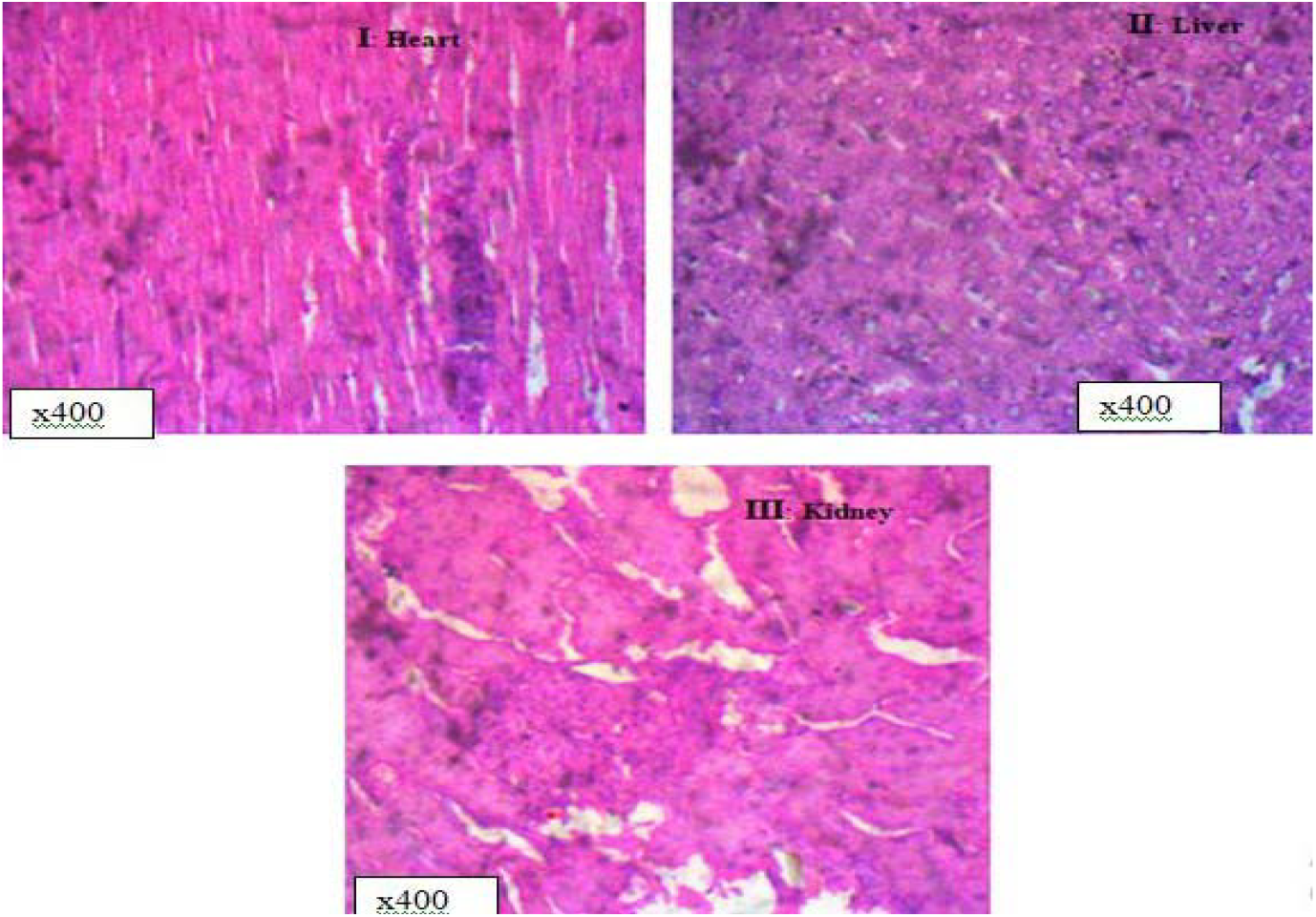
Histology of group 2 animals (negative control) stained with H&E. I) Heart-There is no observable lesion; II) Kidney-There is no observable lesion, III) Liver-Hepatocellular necrosis and inflammation.

**Plate 3.**
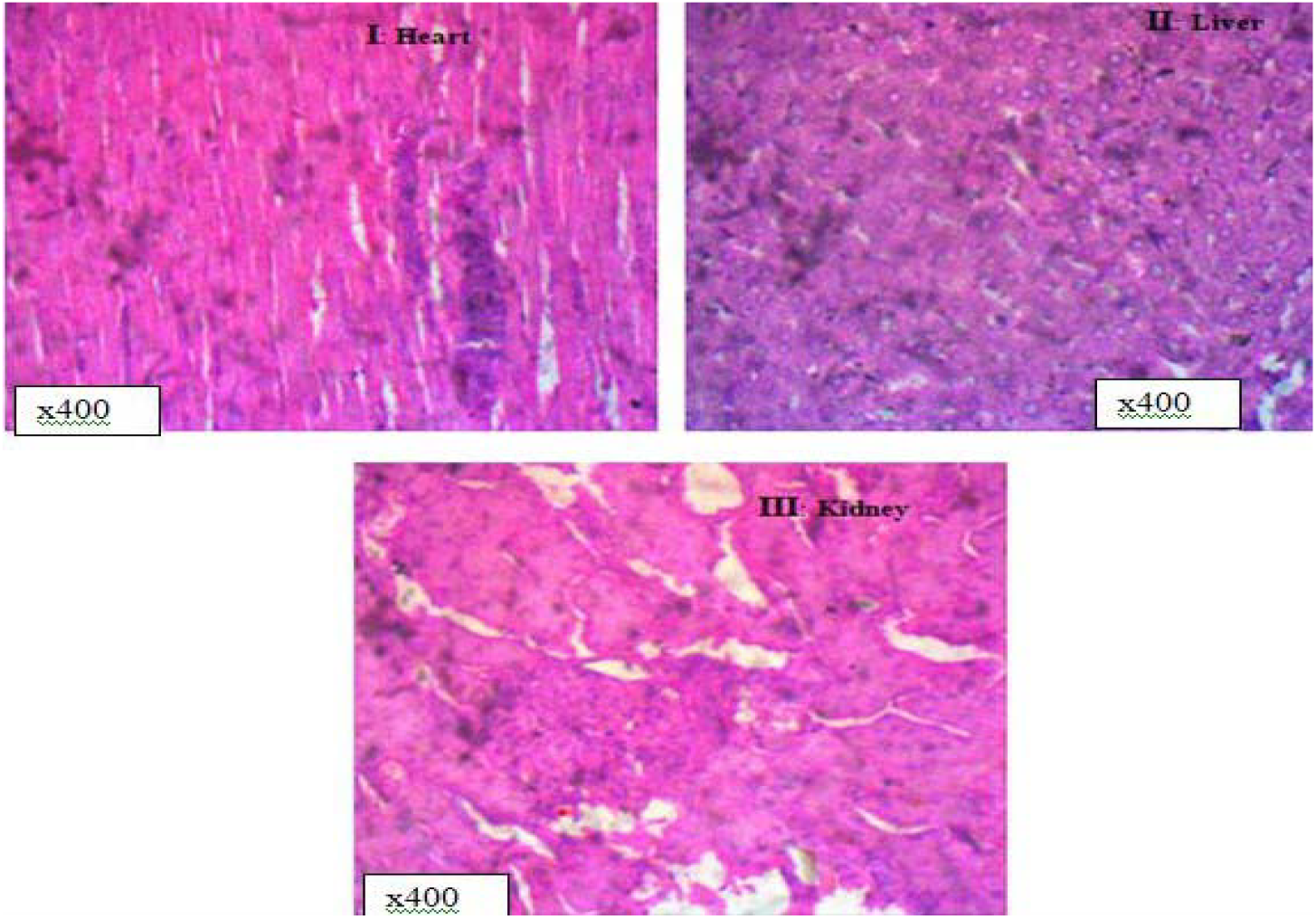
Histology of group 3 animals (treated with standard drugs) stained with H&E. I) Kidney-There is no observable lesion. II) Heart-There is no observable lesion, III) Liver-There is no observable lesion.

**Plate 4.**
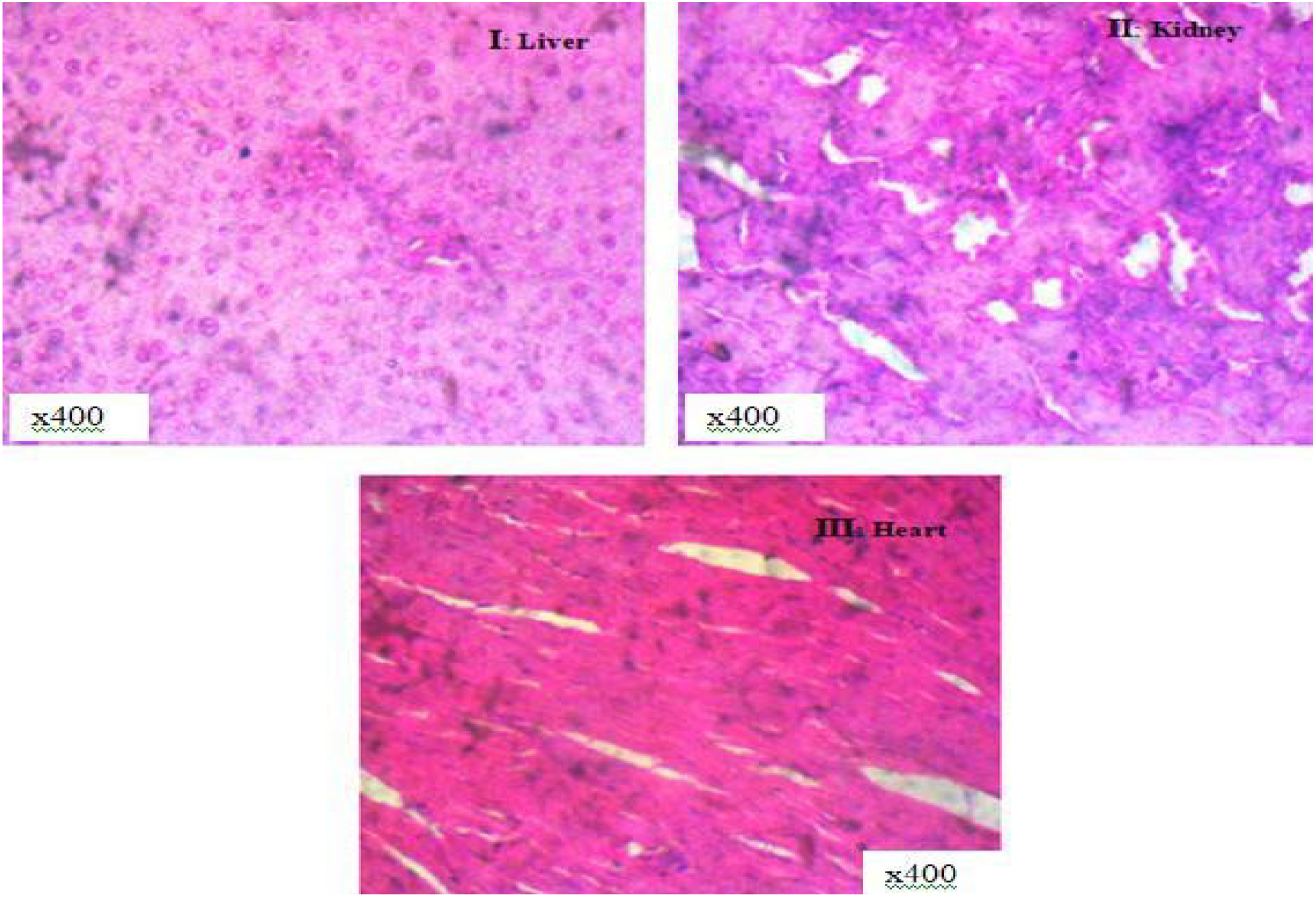
Histology of the effect of seed extract on animals (group 4) stained with H&E. I) Liver-There is no observable lesion, II) Kidney-There is no observable lesion, III) Heart-There is no observable lesion.

**Plate 5.**
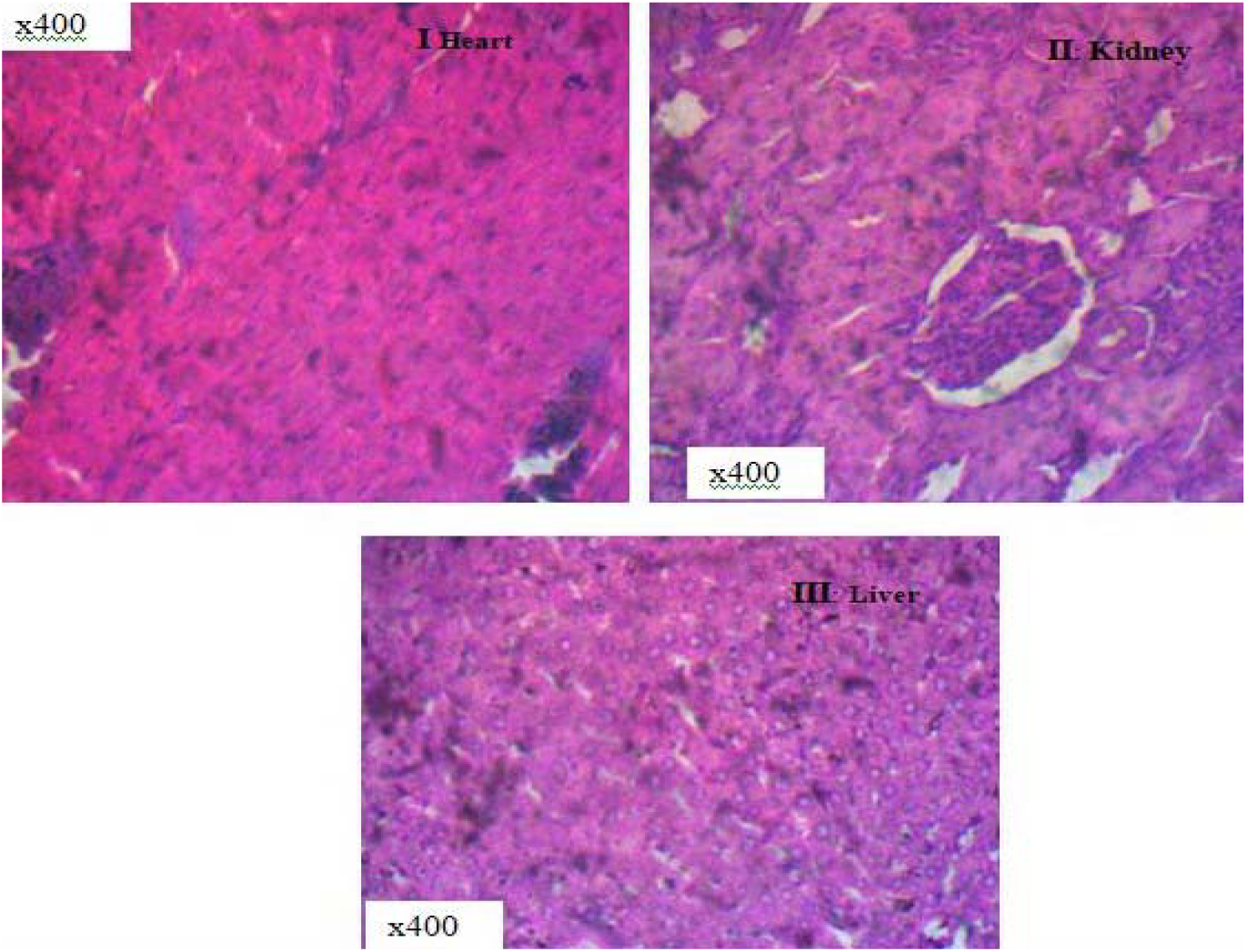
Histology of the effect of seed extract on animals (group 5) (x400). I) Heart-There is no observable lesion, II) Kidney-There is no observable lesion, III) Liver-There is no observable lesion.

**Plate 6.**
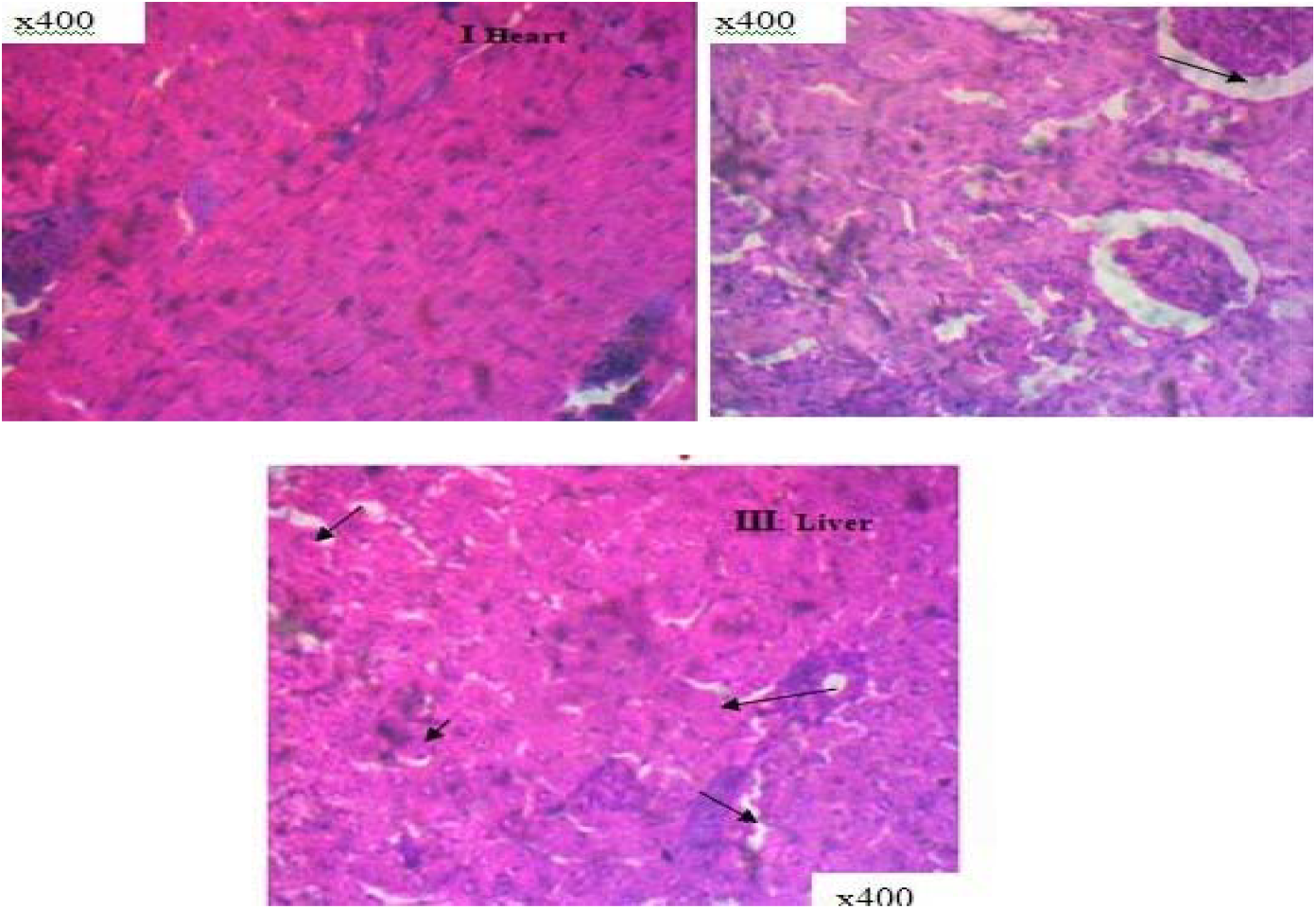
Histology of seed extract’s effect on animals (group 6) stained with H & E stain. I) Heart-There is no observable lesion, II) Kidney: There is moderate atrophy of the glomerular tuft, III) Liver: mild portal inflammation.

**Plate 7.**
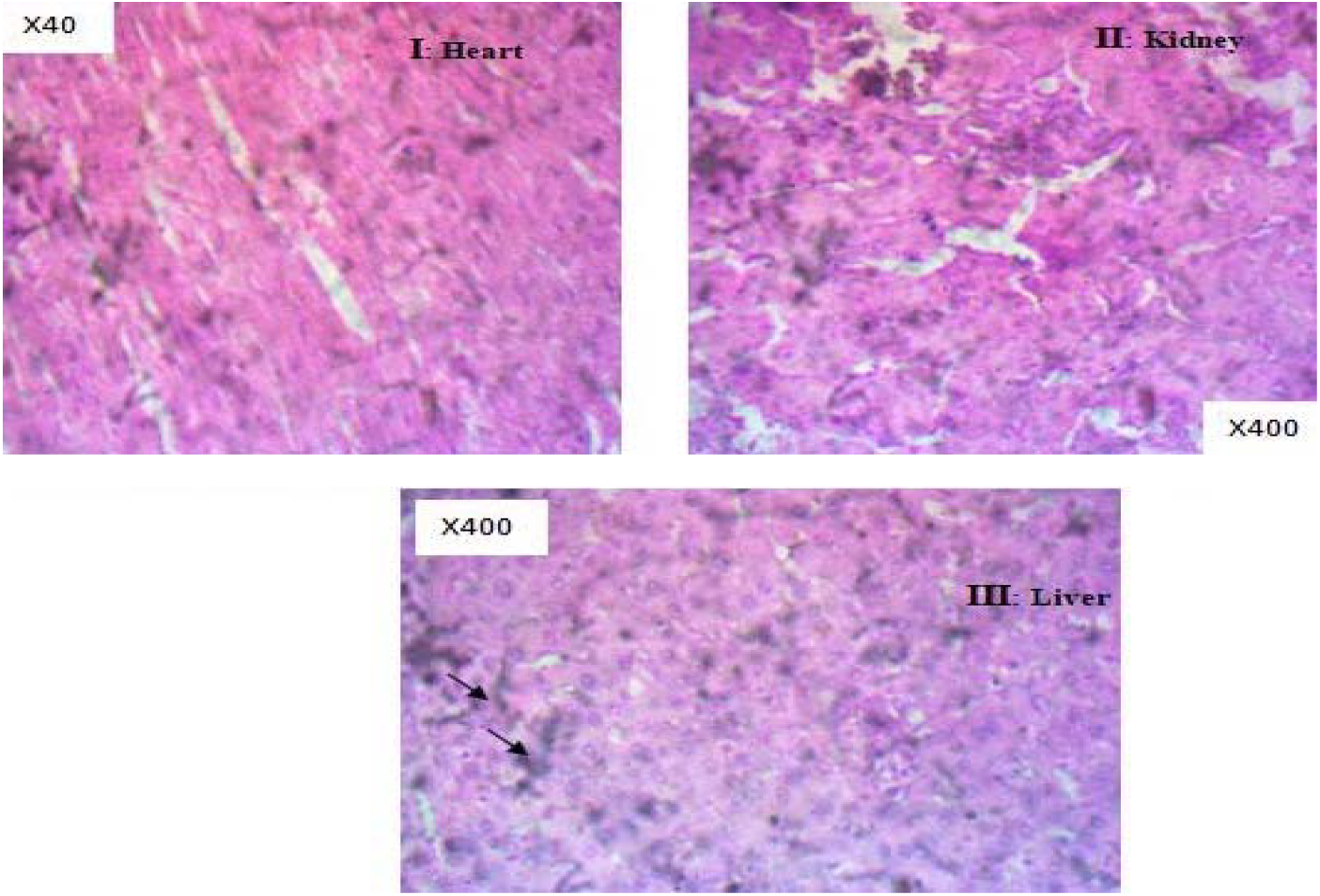
Histology of the effect of seed extracts on H&E-stained animals (group 7). I) Heart-There is no observable lesion HE x400, II) Kidney-There is no observable lesion. HE x400, III) Liver: There is mild portal inflammation.

**Plate 8.**
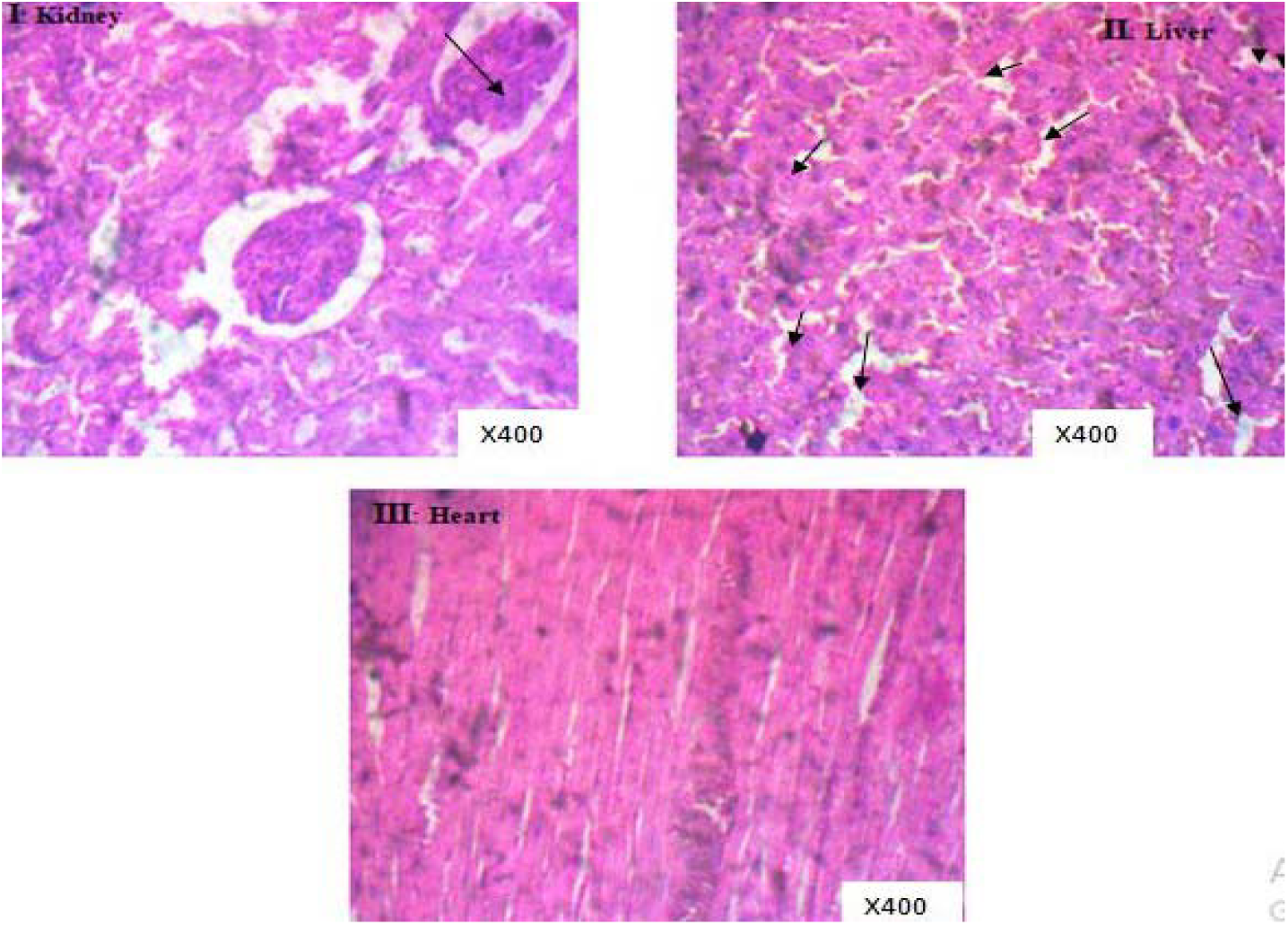
Histology of the effect of the ethanol extract of *C. icaco* seeds on animals (group 8) stained with H&E.I) Kidney-There is moderate atrophy of the glomerular tuft, II) Liver-Hepatocellular necrosis and Kupffer cell hyperplasia, III) There are many observable lesions.

## DISCUSSION

The current study focused on the phytochemical, antioxidant, and antihyperlipademic activities of *C. icaco* seeds. Phytochemical screening revealed the presence of various types of secondary metabolites. The variation in phytochemicals might contribute to the therapeutic basis of *C. icaco* seed extract.

According to the acute toxicity test, the ethanol extract of *C. icaco* seeds caused acute toxicity with an LD50 value *of* 3 807.89 mg/kg, which provides initial evidence of the potential toxic effects of the seed extract. Therefore, the extract of *C*. icaco seeds can potentially cause toxic effects on experimental animals when it is administered at high doses.

In this study, the daily administration of a high-fat diet ad libitum for 28 days resulted in significant weight gain in rats fed a high-fat diet compared to control rats fed standard rat feed. This research corroborates the findings of Al-Otaibi et al. (2020) and Nwankwo et al. (2021), who indicated that animals on a high-fat diet experienced weight gain.

Animals subjected to daily treatment with seed extract (100 to 500 mg/kg) for four weeks exhibited a considerable decrease in weight, comparable to that of animals treated daily with atorvastatin (2 mg/kg), a standard pharmaceutical agent. This result implies that the seed extract can alter the body weight of the animals. The notable reduction in cholesterol, triglyceride, LDL cholesterol, and VLDL cholesterol levels, along with the observed rise in HDL levels at the administered doses (ranging from 100 to 500 mg/kg), substantiates the traditional assertion that the *C. icaco* plant serves as an antihyperlipidemic agent.

Histological evaluations showed that the administration of 300 to 500 mg/kg of the seed extract resulted in organ toxicity comparable to that linked with hepatotoxins and nephrotoxins. This result corroborates the outcomes of the acute toxicity tests conducted on the extract of *C. icaco* seeds. Additionally, the study found that administering the extract at 100 mg/kg and 200 mg/kg did not produce any indications of organ toxicity.

The mild inflammation of the portal area, multifocal necrosis of hepatocytes, and hyperplasia of Kupffer cells observed in the liver, along with the inflammation and moderate atrophy of the = glomerular tuft in the kidney, may be attributed to oxidative stress and liver damage resulting from a high-fat diet, as indicated by Khaleel (2018), Al-Otaibi et al. (2020), and Nwankwo et al. (2021).

The findings suggest that ethanol extracts of *C. icaco* seeds, administered at doses of 100 and 200 mg/kg, were found to be safe and beneficial in enhancing renal and liver functions, as they effectively reversed pathological changes and reinstated the typical structure of both the liver and kidneys. Furthermore, the extract mitigated the detrimental effects that a high-fat diet may have induced. This beneficial effect may be attributed to the antioxidant potentials of the extract, as assessed through DPPH and FRAP evaluations.

## Conclusion

In summary, this research indicated that *C. icaco* lowered elevated cholesterol levels in rats and demonstrated antihyperlipidemic properties, thereby validating its application as a traditional remedy for hyperlipidemia.

## ACKNOWLEDGEMENT

The authors thank the Department of Pharmacognosy, Faculty of Pharmacy, University of Ibadan, and the Department of Pharmacognosy and Traditional Medicine, where the study was carried out, for providing the necessary facilities.

